# Convergent evolution of seasonal camouflage in response to reduced snow cover across the snowshoe hare range

**DOI:** 10.1101/851766

**Authors:** Matthew R. Jones, L. Scott Mills, Jeffrey D. Jensen, Jeffrey M. Good

## Abstract

Determining how different populations adapt to similar environments is fundamental to understanding the limits of adaptation under changing environments. Snowshoe hares (*Lepus americanus*) typically molt into white winter coats to remain camouflaged against snow. In some warmer climates, hares have evolved brown winter camouflage – an adaptation that may spread under climate change. We used extensive range-wide genomic data to 1) resolve broad-scale patterns of population structure and gene flow and 2) investigate the factors shaping the origins and distribution of winter-brown camouflage variation. In coastal Pacific Northwest (PNW) populations, winter-brown camouflage is known to be determined by a recessive haplotype at the *Agouti* pigmentation gene. Our phylogeographic analyses revealed deep structure and limited gene flow between PNW and more northern Boreal populations, where winter-brown camouflage is rare along the range edge. Genome sequencing of a winter-brown snowshoe hare from Alaska shows that it lacks the winter-brown PNW haplotype, reflecting a history of convergent phenotypic evolution. However, the PNW haplotype does occur at low frequency in a winter-white population from Montana, consistent with the spread of a locally deleterious recessive variant that is masked from selection when rare. Simulations show that if annual snow cover dramatically declined in the same population, then the predicted selective increase in frequency of the now beneficial winter-brown *Agouti* allele is likely to be extremely slow due to the same masking effect. Our findings underscore how allelic dominance can shape the geographic extent and rate of convergent adaptation in response to rapidly changing environments.

## Introduction

In response to a shared selection pressure, populations may adapt through the migration of beneficial alleles or through independent mutations that result in the evolution of convergent phenotypes [1–3]. Distinguishing between these scenarios is crucial to understand the capacity of populations to adapt rapidly to environmental change [2,4,5]. If gene flow between populations is sufficiently high, then beneficial variation may spread quickly [6] and potentially allow for rapid adaptive responses to changing environments [7]. However, the spread of adaptive variation may be limited at broad geographic scales, and populations may have to rely on independent standing genetic variation or new mutations to evolve convergent traits. While there is considerable evidence for both adaptation through gene flow and independent mutations in nature [8–14], few empirical studies have examined the specific factors that influence these outcomes in natural populations [2].

When a species range encompasses a mosaic of habitats, the relative probability of adaptation through gene flow versus independent mutation is predicted to be primarily a function of the distance between habitat patches (in units of dispersal distance) and the strength of selection against locally adaptive alleles in intervening habitats [2,15,16]. In general, as distance between patches and the strength of purifying selection in intervening habitats increases, so does the relative probability of adaptation through independent mutations. Adaptation via independent mutations is therefore predicted to be more common in widespread populations where dispersal distance is short relative to the total range size [15]. For example, rock pocket mice (*Chaetodipus intermedius*) have repeatedly evolved melanistic coats across patchy lava flows in the southwestern United States. Although substantial gene flow between adjacent lava flows has likely resulted in the migration of melanic alleles [13], melanism is attributed to different mutations across disparate lava flows [8,17,18]. Theoretical models further predict that the relative probability of adaptation via independent mutations increases rapidly with distance between lava flows (over the scale of tens to hundreds of kilometers) [2], due in large part to strong selection against coat color mismatch [19–21]. Thus, there is a clear tradeoff between dispersal distance and the strength of purifying selection that strongly dictates the probability of adaptation through convergent evolution or gene flow.

The effect of genetic dominance on the probability of convergent evolution has not yet been thoroughly explored [2]. ‘Haldane’s sieve’ [22] predicts that *de novo* dominant mutations enjoy a much greater probability of fixation compared to *de novo* recessive mutations (probability of fixation ≈2*hs*), because rare dominant mutations are visible to selection [23]. As *de novo* beneficial recessives are masked to selection when rare, those that do ultimately reach fixation may spend a longer period of time drifting at low frequency [24]. Likewise, rare recessive migrant alleles are expected to exhibit the same behavior, which may allow more time for mutation to generate ‘competing’ convergent phenotypes upon which selection can act. However, Orr and Betancourt [25] demonstrated that genetic dominance has no effect on the probability of fixation for alleles in mutation-selection balance because recessive alleles have a higher mutation-selection balance frequency. Thus, for relatively high amounts of gene flow, genetic dominance may have little to no effect on the probability of adaptation via migration versus *de novo* mutation. In fact, the masking of low frequency recessive alleles may result in weaker purifying selection in intervening habitats therefore facilitating the spread of adaptive variation through gene flow [2], although this hypothesis remains to be tested.

Snowshoe hares (*Lepus americanus*) are one of at least 21 species that have evolved seasonal molts to white winter coats to maintain camouflage in snowy winter environments. Because color molts are cued by photoperiod, and hence may become mismatched under rapidly changing environments [26,27], seasonal camouflage provides a useful trait to understand adaptation to climate change [26–30]. Reduction in the extent and duration of snow cover is one of the strongest signatures of climate change in the Northern hemisphere [31,32], suggesting that selection should increasingly favor delayed winter-white molts in snowshoe hares to reduce the total duration of coat color mismatch. However, below some annual snow cover threshold, populations are predicted to maintain brown coloration during the winter [28]. Consistent with this, snowshoe hares maintain brown winter camouflage in some temperate environments with reduced snow cover [33], a strategy that should be increasingly favored under climate change [28,29].

Winter-brown snowshoe hares are common in coastal regions of the Pacific Northwest (PNW), where this Mendelian trait is determined by a recessive variant of the *Agouti* pigmentation gene [29]. The locally adaptive *Agouti* variant was introduced into snowshoe hares by hybridization with black-tailed jackrabbits ∼9-18 thousand years ago (kya) and subsequently experienced a local selective sweep within the last few thousand years [34]. Occasional records of winter-brown camouflage also occur in more northern regions along the Pacific coast (e.g., Canada and Alaska) and in eastern North America [28,35]. Although hares are expected to be more or less continuously distributed along suitable Pacific coast environments, there is a deep split and little evidence of recent gene flow between the northern ‘Boreal’ populations and southern ‘PNW’ populations [36,37]. Given this historic population structure, it remains unclear whether the distribution of winter-brown camouflage across populations from disparate parts of the range reflects independent genetic origins (i.e. trait convergence) or spread of the introgressed PNW *Agouti* allele.

Here, we use new and previously published genetic data to investigate the roles of gene flow and mutation in shaping the evolution of winter-brown camouflage across populations of snowshoe hares. We first combined previously published microsatellite (*n*=853 individuals, 8 loci) with new and published whole exome data (*n*=95 individuals) to resolve range-wide patterns of population history and gene flow in snowshoe hares, which provides crucial context for understanding the historical spread and adaptive potential for winter-brown camouflage to climate change. We then generated whole genome sequence (WGS) data of a winter-brown Alaska (AK) hare to test whether winter-brown camouflage in Boreal snowshoe hares arose independently from PNW populations. Next, to understand the geographic limits of the recessive PNW *Agouti* allele, we used pooled WGS data to estimate its frequency in a winter-white population from Montana (MT), approximately 600 km in a straight-line from the closest winter-brown PNW population. Finally, we used both theoretical predictions and simulations to understand the factors influencing the geographic scope of the PNW *Agouti* allele and its potential to contribute to rapid adaptation in response to warming climates.

## Methods

### Samples and genomic data generation

To resolve patterns of range-wide population structure, we performed targeted whole exome enrichment of 12 snowshoe hares previously sampled from 12 localities across Canada, Alaska, and the eastern United States and three hares from the southern Rocky Mountains (representing the ‘Boreal’ and ‘Rockies’ lineages as defined by Cheng et al. [36]). The targeted whole exome capture was designed by Jones et al. [29] to enrich for ∼99% of genic regions annotated in the European rabbit (*Oryctolagus cuniculus*) genome (61.7-Mb spanning 213,164 intervals; ∼25-Mb protein-coding exons, ∼28-Mb untranslated region, ∼9-Mb intronic or intergenic). We followed the library preparation protocols outlined in Jones et al. [29] and sequenced libraries on one lane of Illumina HiSeq2500 with paired-end 100 bp reads (HudsonAlpha Institute for Biotechnology; Huntsville, AL). Exome sequences for Boreal and Rockies snowshoe hares were combined with published whole exome data [29] from 80 PNW snowshoe hares (*n*=95 total, Table S1), including a monomorphic winter-brown population from southeast British Columbia (BC1, *n*=14), a monomorphic winter-white population from Seeley Lake area in western MT (*n*=14), and two polymorphic coat color populations from Oregon (OR, *n*=26; two localities) and Washington (WA, *n*=26) [29].

To resolve the historical spread of the winter-brown PNW *Agouti* allele, we performed whole genome sequencing of a winter-brown snowshoe hare museum specimen (AK) from southwest Alaska first noted by Link Olson and later obtained through loan from University of Alaska Museum of the North (UAM 116170, collected on 4 January 2013). We also performed whole genome sequencing on a DNA pool of 81 snowshoe hares from two localities in Glacier National Park, MT (Table S1). We extracted genomic DNA from muscle tissue of the AK hare sample following the Qiagen DNeasy Blood and Tissue kit protocol (Qiagen, Valencia, CA). For the pooled MT whole genome sequencing, we pooled previously extracted DNA samples [36] in approximately equimolar quantities based on Qubit concentrations (Invitrogen Qubit Quantitation system LTI). We then prepared genomic libraries for all samples following the KAPA Hyper prep kit manufacturer’s protocol. We sheared genomic DNA to ∼300 bp using a Covaris E220evolution ultrasonicator and performed a stringent size selection using a custom-prepared carboxyl-coated magnetic bead mix [38]. We determined indexing PCR cycle number for each library with quantitative PCR (qPCR) on a Stratagene Mx3000P thermocycler (Applied Biosystems) using a DyNAmo Flash SYBR Green qPCR kit (Thermo Fisher Scientific). Final libraries were size-selected again with carboxyl-coated magnetic beads, quantified with a Qubit (Thermo Fisher Scientific), and pooled for sequencing by Novogene (Novogene Corporation Ltd.; Davis, CA) on two lanes of Illumina HiSeq4000 using paired-end 150bp reads. The whole genome sequence from the AK hare was combined with previously published whole genome sequencing of five black-tailed jackrabbits, three winter-brown snowshoe hares from OR, WA, and BC1, and three winter white snowshoe hares from MT, a hare from Utah (UT), and a hare from Pennsylvania (PA, Table S1) [29].

### Read processing and variant calling

For all raw Illumina sequence data, we trimmed adapters and low-quality bases (mean phred-scaled quality score <15 across 4 bp window) using Trimmomatic v0.35 [39] and merged paired-end reads overlapping more than 10 bp and with lower than 10% mismatched bases using FLASH2 [40].

Whole genome sequence data were mapped to either a snowshoe hare or black-tailed jackrabbit pseudoreference (see Jones et al. [29] for details) using default settings in BWA-MEM v0.7.12 [41]. We used *PicardTools* to remove duplicate reads with the MarkDuplicates function and assigned read group information with the AddOrReplaceReadGroups function. Using GATK v3.4.046 [42], we then identified poorly aligned genomic regions with RealignerTargetCreator and locally realigned sequence data in these regions with IndelRealigner. We performed population-level multi-sample variant calling using default settings with the GATK UnifiedGenotyper. Here, we called variants separately for each previously defined snowshoe hare population genetic cluster (i.e., Boreal, Rockies, BC1, MT, OR, and WA) and for black-tailed jackrabbits. We performed variant filtration in VCFtools v0.1.14 [43]. For whole exome and whole genome data, we filtered genotypes with individual coverage <5× or >70× or with a phred-scaled quality score <30. Additionally, we removed all indel variants and filtered SNPs with a phred-scaled quality score <30, Hardy-Weinberg *P*<0.001. We required that sites have no missing data across individuals.

### Range-wide population genetic structure and gene flow

We inferred a maximum likelihood tree with a general time reversible model in RAxML v8.2.8 [44] using a subset of the concatenated snowshoe hare exome data (*n*=12 Boreal hares, *n*=3 Rockies hares, *n*=12 PNW hares; 21,167,932 total sites) with European rabbit (*Oryctolagus cuniculus*) as the outgroup. Using this maximum likelihood tree as a starting tree, we estimated a maximum clade credibility tree and node ages with a constant population size coalescent model in BEAST 2 [45]. We assumed a strict molecular clock and an HKY substitution model using empirical base frequencies. We specified default priors for the kappa parameter, gamma shape parameter, and population size parameter and used a gamma distribution (alpha=0.0344, beta=1) as a prior for the clock rate parameter. We ran the MCMC for 5 million steps and calibrated divergence times using a log-normal distribution for the *Oryctolagus*-*Lepus* node age with a median of 11.8 million years (95% prior density: 9.8–14.3) [46]. We also performed a species-tree analysis using gene trees in ASTRAL v5.6.3 [47]. Gene trees were generated across 200 kb windows (excluding windows wither fewer than 500 sequenced bases) using RAxML with a GTR+gamma model and rapid bootstrap analysis (-f a -# 10). We collapse nodes on gene trees with low bootstrap support values (<=10) and performed ASTRAL analyses on the collapsed gene trees using default settings.

To test for signatures of population structure and gene flow among lineages, we performed a population cluster analysis in ADMIXTURE v1.3.0 [48]. We tested *K* values from 1-10 (representing the number of population clusters) and selected the *K* value with the lowest cross-validation error. We also estimated range-wide effective migration and diversity surfaces with EEMS [49] using extensive microsatellite data (*n*=853 individuals, 8 loci) generated by Cheng et al. [36]. Varying the number of demes (50, 100, and 200) had little effect on estimates of effective rates of migration and diversity, therefore we only report results for 200 demes. We used default hyper-parameter values and tuned the proposal variances such that proposals were accepted approximately 30% of the time. We ran EEMS for 2 million iterations with a burn-in of 1 million iterations and thinning iteration of 9999. Runs produced strong correlations between observed and expected genetic dissimilarity both within and between demes, indicating good model fit.

### Geographic distribution of the PNW *Agouti* allele

The winter-brown AK snowshoe hare was collected approximately 3000 km (via the Pacific coast) from Vancouver, BC, the northern limit of winter-brown PNW hare populations [29]. To test whether the introgressed PNW *Agouti* allele has seeded winter-brown camouflage in southwest AK, we generated a tree of the PNW *Agouti* haplotype region from our whole genome sequence data of winter-brown and winter-white snowshoe hares (*n*=7) and black-tailed jackrabbits (*n*=5). We defined the PNW *Agouti* haplotype region (chr4:5,340,275 – 5,633,020, oryCun2 coordinates) based on the location of the introgressed black-tailed jackrabbit tract identified by Jones et al. [29] using a hidden Markov model. We then inferred a maximum likelihood phylogeny for this interval using RAxML [44] as above. Node support values were generated from 1000 replicate bootstrap runs.

We also estimated the frequency of the PNW *Agouti* haplotype among pooled individuals from two localities in MT (*n*=81) that are 575 km and 627 km from the polymorphic sampling locality in WA, where winter-brown camouflage is relatively common. We used PoPoolation2 [50] to calculate the frequency of the winter brown-associated alleles at 555 sites across the introgressed *Agouti* haplotype (chr4: 5,340,275 – 5,633,020). These positions were previously shown to be strongly associated with coat color based on a likelihood ratio test of allele frequency differences between winter-brown and winter-white individuals from low coverage whole genome sequence data [29]. We summed the counts of winter-white and winter-brown alleles across all positions to estimate both the mean and standard deviation of winter-brown allele frequency. We excluded seven positions with unusually high frequencies of the winter-brown allele (∼45-100%) supported by reads that did not otherwise carry brown-associated alleles (i.e., incongruent haplotype information), as these likely reflect sequencing errors.

### Probability of convergent adaptation through independent mutation

To compare the observed spread of the PNW *Agouti* allele to theoretical expectations, we estimated the relative probability of adaptation via independent mutations with distance from a focal habitat patch using the model developed by Ralph and Coop [2] (equation 12) under a range of conditions. Since the relative probability of adaptation through migration or *de novo* mutation depends strongly on the rate at which mutations generate convergent phenotypes [2], we tested a range of plausible mutation rates. A previous study using over 7 million house mice estimated a mean rate of spontaneous visible coat color mutations of 11e^-6^ per locus/gamete [51], which is ∼1930× higher than the average genome-wide germline per site mutation rate (5.7e^-9^ per site/gamete) [52]. Given a genome-wide mutation rate of 2.35e^-9^ per site/generation for rabbits [53], we tested a high mutation rate of *μ*=4.54e^-6^ (i.e., the overall expected rate of visible coat color mutations) and a low mutation rate of *μ*=4.54e^-8^ (i.e., assuming only 1% of coat color mutations would lead to brown winter camouflage). We also tested how strong (*s*_*m*_=0.01), moderate (*s*_*m*_=0.001), or weak (*s*_*m*_=0.0001) purifying selection against the winter-brown allele in intervening (i.e., snow-covered) habitats influences the probability of adaptation through independent mutations. We assumed a relatively high mean dispersal distance of 2 km [54], which is likely to produce a conservative estimate of the probability of adaptation via independent mutation. We also assumed that the width of the secondary habitat patch was relatively small (*w*=1 or ∼6-45 km, depending on *s*_*m*_), which is consistent with observations of winter-brown camouflage at low frequency along portions of the range edge. Finally, we assumed *d*=1 (dimension of the habitat), *C*=1, and *γ*=0.5.

### Simulations of selection on migrant alleles with genetic dominance

Theory predicts that allelic dominance should not influence a mutation’s probability of fixation under positive selection if it starts in mutation/migration-selection balance [25]. Nonetheless, rare recessive mutations may increase in frequency more slowly relative to rare dominant mutations because they are initially invisible to selection, allowing more opportunity for competition from convergent phenotypes arising through independent mutations. We used simulations to test how the genetic dominance of alleles at equivalent mutation-selection equilibrium frequencies influences the probability and rate of adaptation to new habitats when environmental conditions change. In SLiM 3.0 [55] we performed 100 simulations of the MT population (estimated *N*_*e*_=245430) [34] with a positively selected recessive mutation (*s*=0.01, *h*=0) starting at the inferred frequency of the PNW *Agouti* allele in MT (*p*). We assumed that *p* reflects the equilibrium frequency (*x*) between the migration rate of the allele into the environment (*m*) and the selection coefficient against the allele (*s*), which is given as 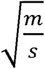 for a recessive mutation. Given that *x* for a dominant mutation is simply 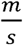, we calculated *x* of an equivalent dominant mutation (same values of *m* and *s*) as simple *p*^2^. We then repeated these simulations for a positively selected (*s*=0.01), completely dominant allele (*h*=1) starting at its expected mutation-selection balance frequency. For each simulation we tracked the frequency of the selected allele until it was either fixed or lost. To validate these simulations, we compared the probabilities of allelic fixation from simulations to fixation probabilities analytically derived by Kimura [56]. To test for significant differences in fixation probabilities and rates of adaptation between recessive and dominant mutations we used Student’s t-tests in R [57].

## Results

### Snowshoe hare phylogeography

We combined previously generated whole exome data for 80 PNW snowshoe hares (mean coverage 21× ± 7.6 per individual) with new whole exomes of 15 hares across the range sequenced to a mean coverage of 20.2× ± 8.2 (Fig. 1A). Exome-wide phylogenetic analyses show three broad phylogeographic groups of snowshoe hares (Fig. 1B) with unambiguous ASTRAL support scores using 6582 gene trees, consistent with previous studies [36,37]. However, ∼12-25% of SNPs within phylogeographic groups appear to be shared across some population boundaries (Table 1), indicating some amount of shared ancestral variation or gene flow. We estimated that snowshoe hares from Canada, Alaska, and the eastern United States— representing the Boreal lineage—diverged approximately 97.2 kya (95% posterior density: 77.4-120.8 kya) from PNW and Rockies snowshoe hares (Fig. 1B). PNW and Rockies hare populations are estimated to have split approximately 47.1 kya (95% posterior density: 37.2-58.2 kya).

**Table 1.** Counts of the number of private (e.g., Boreal/Boreal) versus shared SNPs (e.g., Rockies/Boreal) for each major snowshoe hare clade. Individuals from WA represent the PNW.

|  | Boreal<br>(*N*=120978 SNPs) | Rockies<br>(*N*=158207 SNPs) | PNW<br>(*N*=275614 SNPs) |
| --- | --- | --- | --- |
| Boreal | 106051 | 3760 | 5820 |
| Rockies & PNW | 5347 | - | - |
| Rockies | - | 117863 | 31237 |
| PNW | - | - | 233210 |

**Figure 1.**
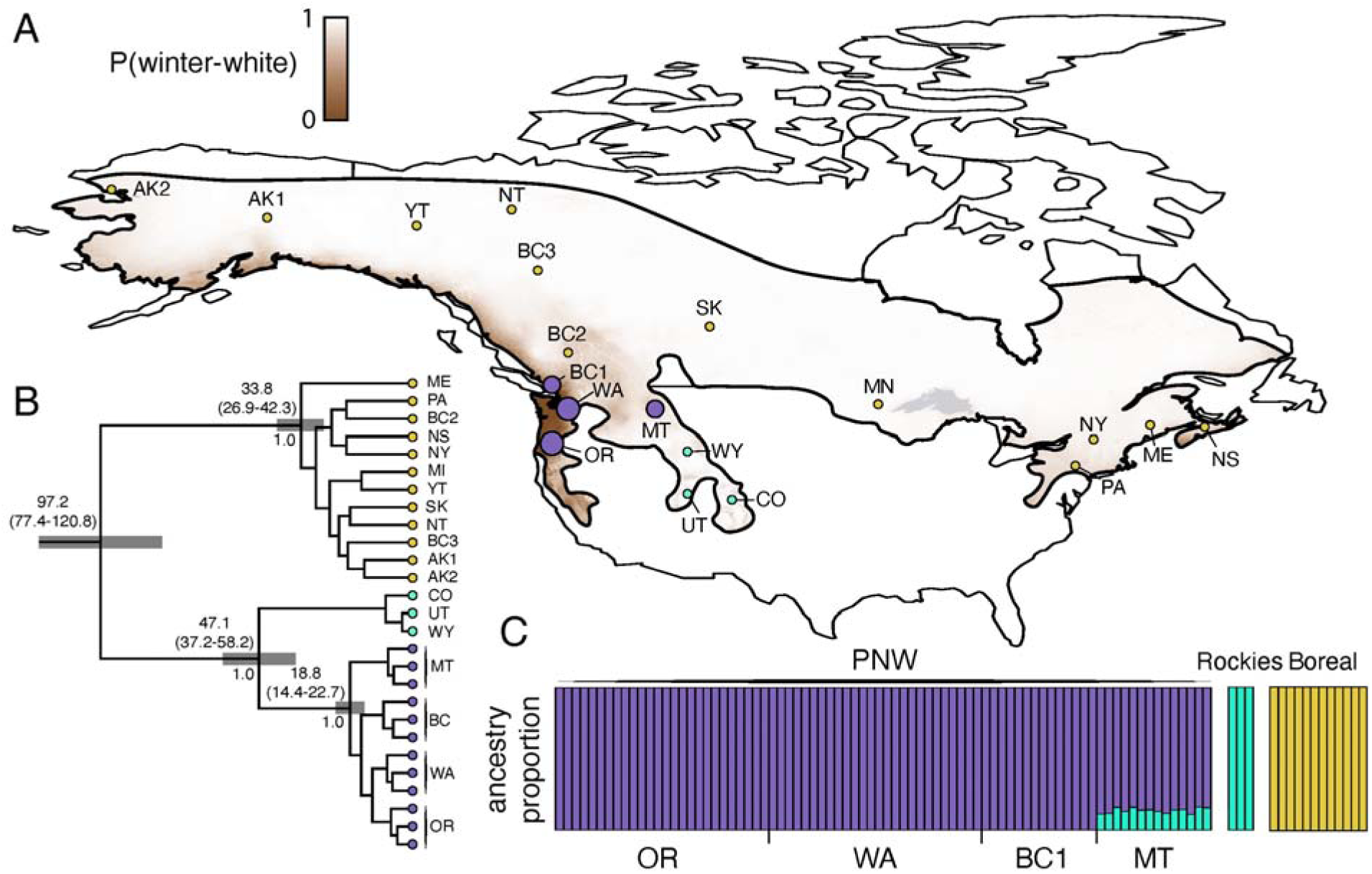
(**A**) Range-wide phylogeography of snowshoe hares based on whole exome sequences. The snowshoe hare range is colored brown to gray according to the modelled probability of winter-brown camouflage from Mills et al. (2018). Sizes of sampling locality circles are scaled to sample size and are colored according to their population assignment (see C). (**B**) A maximum credibility phylogenetic tree estimated with BEAST 2 [45]. All nodes have posterior probabilities >99%. Each major node shows the median node age in thousands of years (95% posterior density in parentheses and gray rectangles) and the ASTRAL support score. (**C**) The Admixture plot shows the proportion of ancestry across samples based on a *K*=3 clustering, which had the lowest cross validation error.

Population structure analyses in Admixture also found strongest support for three population clusters, although MT snowshoe hares (in the PNW lineage) showed 11-16% of their ancestry assigned to the Rockies lineage, indicating ongoing gene flow or continuous genetic structure (Fig. 1C). The Admixture analysis also indicated little apparent gene flow between the Boreal lineage and PNW or Rockies lineages. Likewise, microsatellite-based estimates of effective migration surfaces revealed that effective migration rates are approximately 100-fold lower than the range-wide average near the apparent zone of contact between PNW and Boreal lineages in western North America (Fig. 2A). We also estimated relatively low effective migration rates across the southwestern edge of snowshoe hare range (western US; log(*m*) ≈-1, or 10-fold lower than the mean). In contrast, effective migration rates in the northwest part of the range (Alaska and western Canada) appeared relatively high (∼10-fold higher than the mean). Effective diversity surfaces show that the highest relative genetic diversity occurs in the eastern extent (Boreal lineage) and southwestern extent (PNW lineage) of the snowshoe hare range (Fig. 2B), consistent with previous studies suggesting that these regions were likely glacial refugia [36]. Relative genetic diversity gradually decreased moving from east to west across the Boreal range and the lowest genetic diversity was observed in the Rocky Mountain lineage, distributed across the western US.

**Figure 2.**
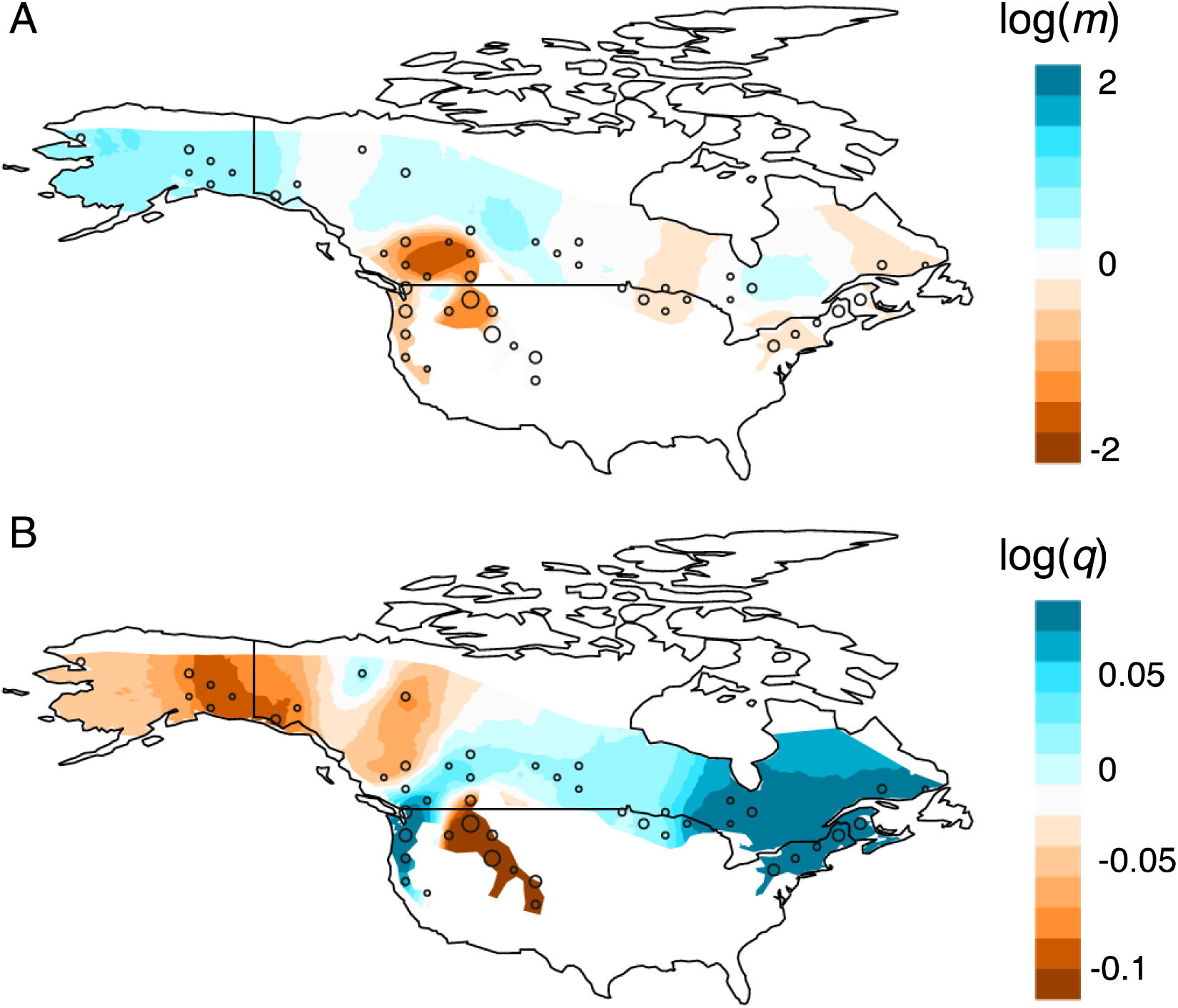
(**A**) Effective migration rates for snowshoe hares inferred from range-wide microsatellite data set (853 individuals, 8 loci) from Cheng et al. (2014). The sizes of circles are scaled to the number of samples collected at that location. Effective migration rate is measured as the rate of decay in genetic similarity of individuals across space. Regions that are colored white are characterized by isolation-by-distance while regions that are colored blue or red have higher or lower effective migration, respectively. (B) Effective diversity rates based on the same microsatellite data set. Here effective diversity rates measure the genetic dissimilarity between individuals in the same deme, where blue regions have higher than average diversity and red regions have lower than average diversity.

### Convergent evolution and the distribution of winter-brown camouflage variation

Under a single origin of winter brown camouflage, we would expect the winter-brown AK individual to nest within the black-tailed jackrabbit clade with the other winter-brown snowshoe hares from the PNW. However, our phylogenetic analysis indicates that the winter-brown AK individual unambiguously groups with a winter-white Boreal hare from PA (100% bootstrap support), and more broadly with other winter-white hares across the range (Fig. 3). These results demonstrate that the winter-brown phenotype of this AK hare is caused by mutations that are independent from the introgressed *Agouti* haplotype that encodes winter-brown camouflage in the PNW, approximately 3000 km away. Determining the genetic basis of this independent origin of winter brown camouflage will likely require an independent genetic association study in these populations.

**Figure 3.**
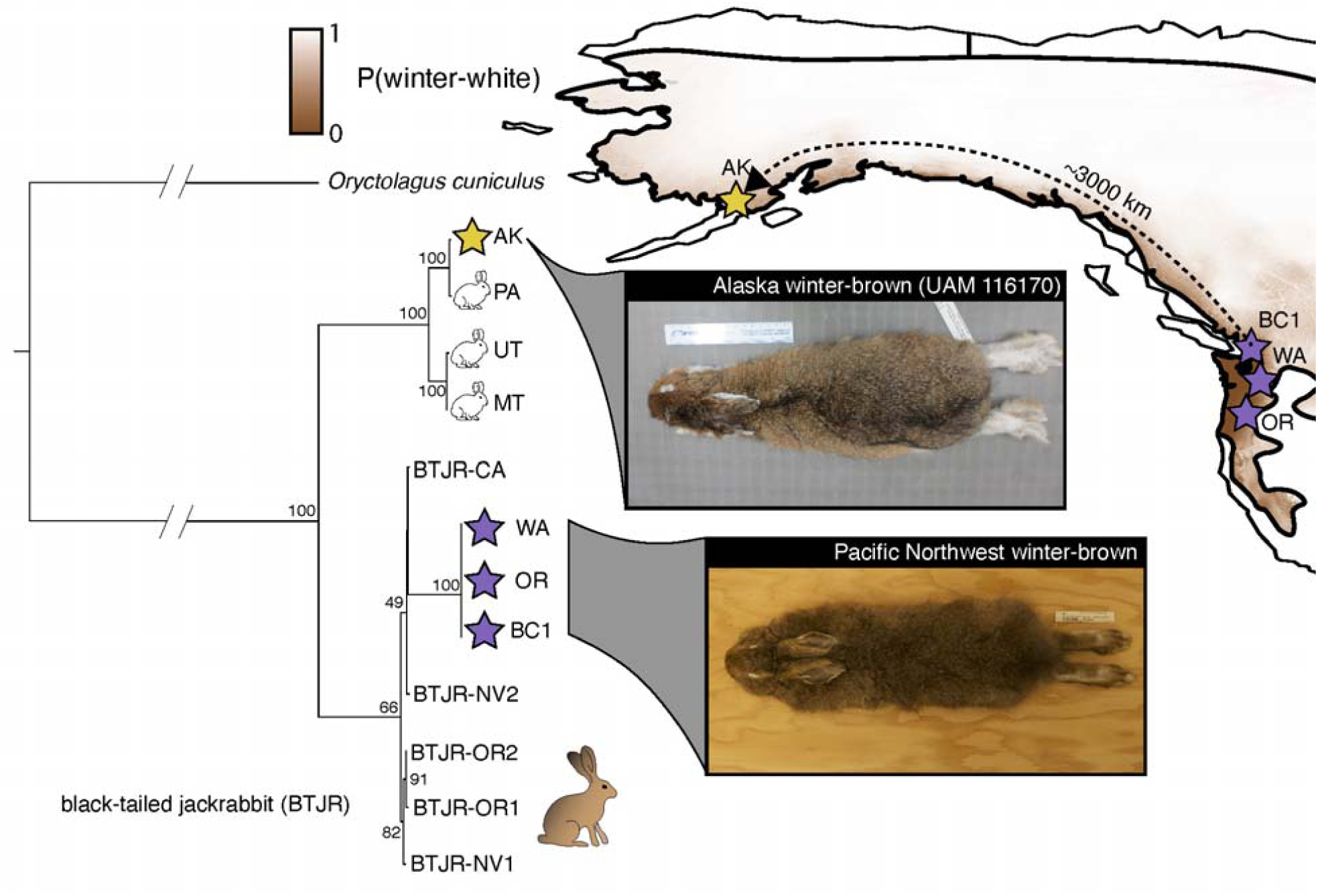
A maximum clade credibility tree of the introgressed *Agouti* locus (∼293 kb) based on whole genome sequencing of black-tailed jackrabbits (BTJR; black circles) and snowshoe hares. Values indicate node support based on bootstrapping. Colored stars indicate winter-brown snowshoe hares from Alaska (yellow, UAM 116170 pictured top) or the Pacific Northwest (purple, pictured bottom).

Pooled whole genome sequencing detected that the recessive winter-brown *Agouti* allele occurs at an estimated frequency of 1.24% (± 0.01%) across predominately winter-white MT localities ∼600 km from the polymorphic zone. Interestingly, long-term live-trapping and radiotelemetry-based field work from this region has resulted in a single observation of a winter-brown hare in approximately 300 winter hare observations (0.33%; Scott Mills pers. obs.), although this observed frequency is slightly higher than the Hardy-Weinberg expectation under the estimated frequency of the winter-brown *Agouti* allele (*p*^*2*^=0.015%, binomial test *p*-value=0.044).

To understand the factors that have shaped the geographic distribution of the PNW *Agouti* allele, we modelled the relative probability of adaptation through independent mutation with distance from a focal habitat using the theoretical framework of Ralph and Coop [2]. The change in relative probability of adaptation through independent mutation depended strongly on both the mutation rate and selection coefficient against the allele in intervening habitats (Fig. 4). Stronger selection coefficients produced a more rapid shift in the convergence probability with distance relative to weaker selection (Fig. 4). For instance, under a high mutation rate to the winter-brown phenotype (*μ*=4.54e^-6^) and strong purifying selection (*s*_*m*_=0.01) the probability of independent adaptation transitioned sharply from *P*=0.1 at 78 km to *P*=0.9 at 140 km from a focal habitat. Under a low mutation rate to the winter-brown phenotype (*μ*=4.54e^-8^) and weak selection (*s*_*m*_=0.0001), independent adaptation was much less likely at close distances, as expected, and the probability increased more gradually (*P*=0.1 at 778 km to *P*=0.9 at 1400 km).

**Figure 4.**
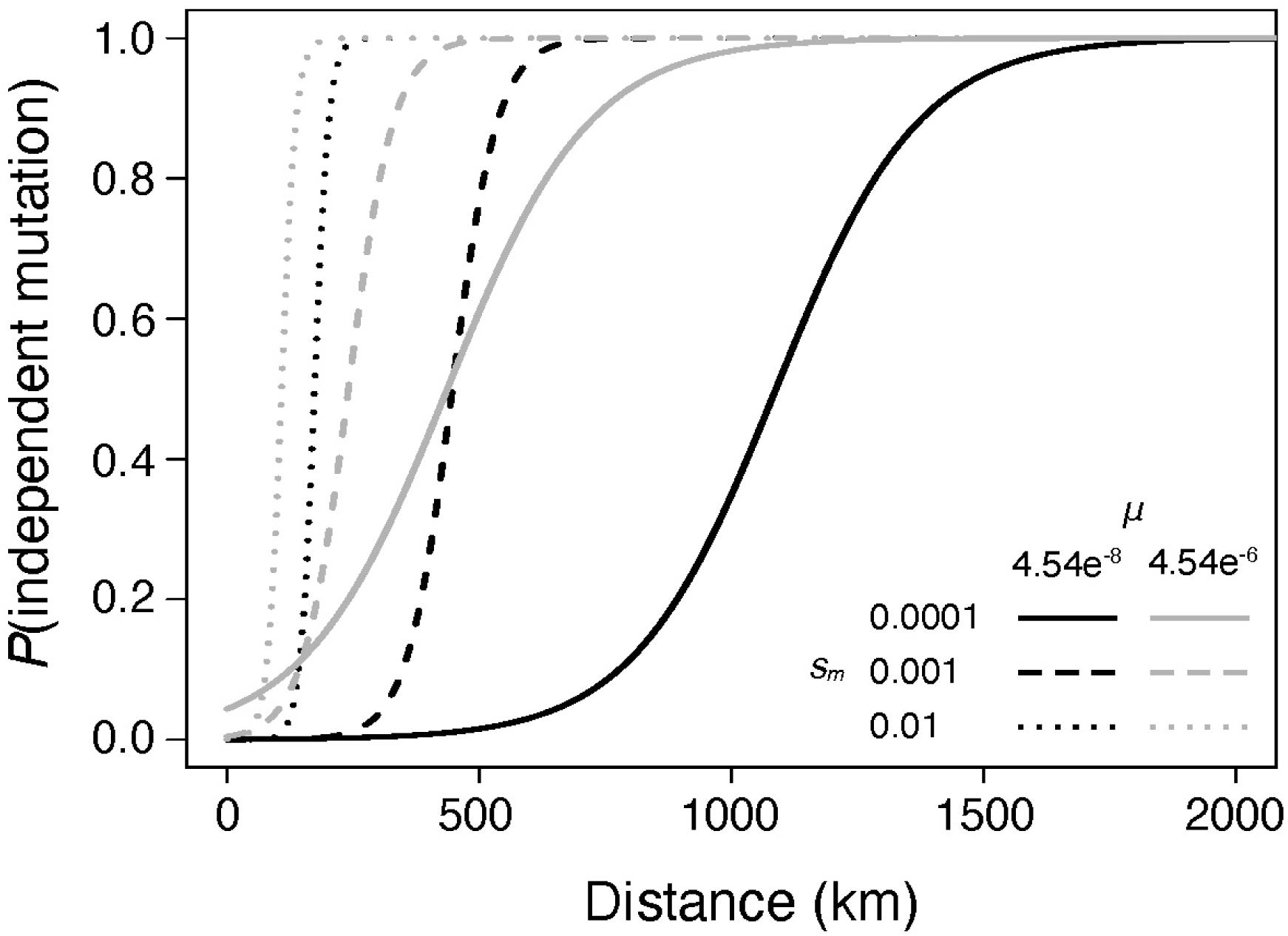
The probability of adaptation through independent mutations in snowshoe hares as a function of distance in kilometers from a focal habitat patch harboring a locally adaptive variant. The probability of independent mutation is calculated according to equation 12 in Ralph and Coop (2015). Here we varied the mutation rate to the winter-brown phenotype (*μ*; black=4.54e^-8^, gray=4.54e^-6^) and the negative selection coefficient in intervening habitats (*s*_*m*_; solid line=0.0001, dashed line=0.001, dotted line=0.01).

### Migration-selection balance

Assuming an environmental shift towards less snow results in strong positive selection (*s*=0.01) on standing variation of the recessive (winter-brown) *Agouti* allele in MT, we used simulations to estimate that the corresponding probability of fixation would be ∼81% (95% confidence intervals: 72.2-87.5%; Fig. 5A), which is consistent with an analytically-derived probability of 78.1%. For an equivalent dominant mutation starting in mutation-selection balance frequency (0.015%) the simulated probability of fixation was approximately 77% using simulations (95% confidence intervals: 67.8-84.2%) or 77.9% using the Kimura [56] equation. Neither estimate was significantly different from the fixation probability of a recessive mutation (*p*=0.48, two-tailed test of equality of proportions). However, we observed striking differences in the rates of increase, conditional on fixation, of dominant and recessive mutations following a sudden environmental change (Fig. 5B). The mean time to fixation was significantly faster (*p*<2.2e-16, Student’s T-test) for a recessive mutation (9645 generations, SD=3609 generations) compared to a dominant mutation (19445 generations, SD=7273 generations, *p*<2.2e^-16^). However, as expected, we see that initial rates of allele frequency change and phenotypic adaptation are considerably faster for a positively selected dominant mutation (Fig. 5B). For instance, the mean time for the selected phenotype determined by the recessive mutation to reach a frequency of 75% (*p*=0.866) was 8837 generations (SD=3630 generations), compared to 1007 generations (SD=116 generations) if determined by the dominant mutation. The striking difference in the rate at which beneficial dominant versus recessive alleles contribute to adaptation is maintained even up to the point at which the selected phenotype reaches a frequency of 99%, which takes approximately 9262 generations (SD=3624 generations) for a recessive mutation and only 2030 generations for dominant mutation (SD=141 generations).

**Figure 5.**
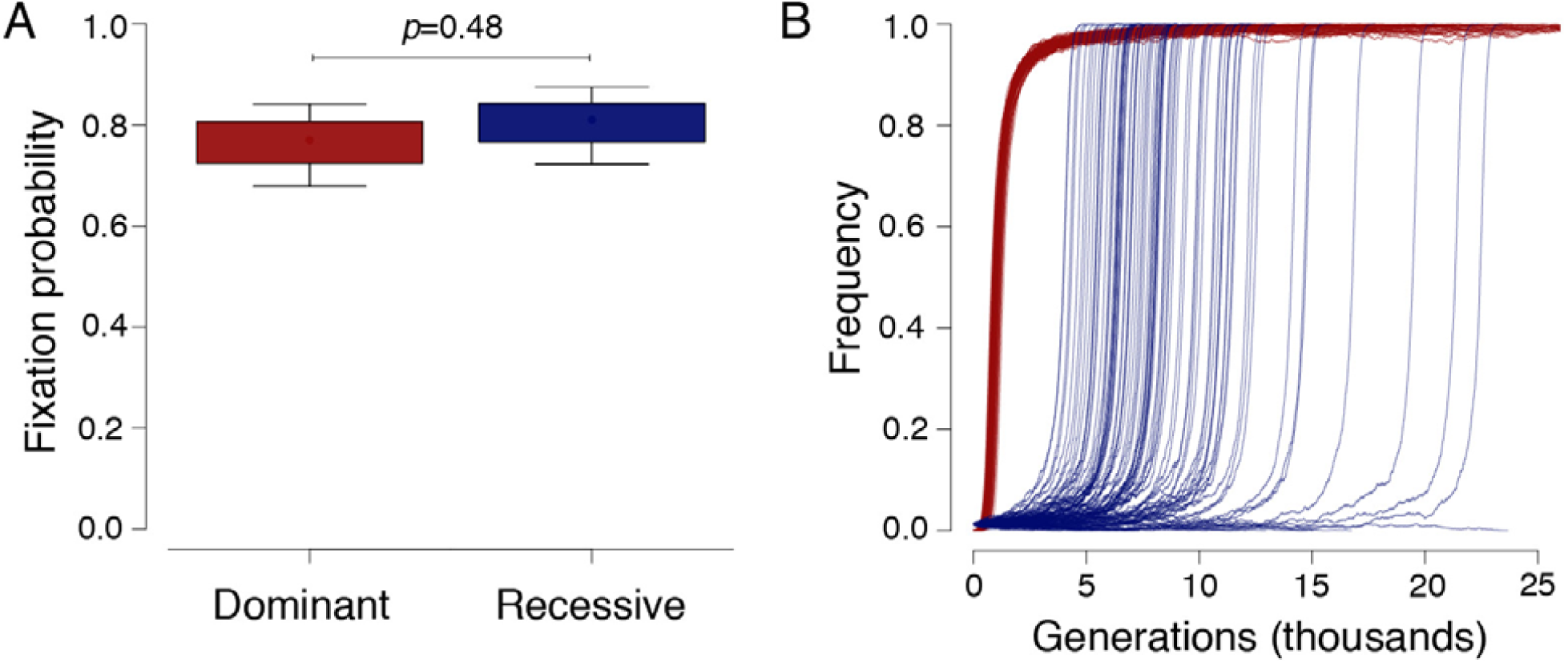
(**A**) The simulated probability of fixation of a completely dominant (red, mean=77%, *N*=100) or recessive (blue, mean=81%, *N*=100) mutation experiencing positive selection and starting in migration-selection balance frequency (0.015% for dominant, 1.24% for recessive). (**B**) The simulated allele frequency trajectories of the same dominant (red) and recessive (blue) mutations starting in migration-selection balance.

## Discussion

A growing number of studies have found evidence for convergent adaptation within and between species [8–12,18,58,59], although we often lack an understanding of the forces that determine whether local adaptation occurs through independent *de novo* mutations or migration of pre-existing alleles from other populations [2]. Our study provides rare empirical insights into how gene flow, mutation, allelic dominance, and selection interact to shape the spatial scale and pace of local adaptation to new or changing environments.

Understanding the potential for adaptive variation to spread between populations across large spatial scales requires basic insights into range-wide patterns of population structure and gene flow. Winter-brown camouflage in snowshoe hares occurs across only ∼5% of the total range but is broadly-distributed from the western edge of the range along the Pacific coast to the eastern extent of the range in Pennsylvania [28,33,35]. Previous phylogeographic studies based on limited genetic data suggest that this distribution spans a deep phylogenetic boundary between Boreal and PNW lineages (∼2 Myr divergence) [36,37] with little evidence for gene flow. With whole exomes (*n*=95), our phylogenetic analysis also supports a deep split but suggests a substantially more recent divergence time between Boreal and Rockies/PNW lineages than previously estimated (∼97 kya; Fig. 1B). Our more recent estimates may be attributable to the increased power of whole exomes or heavy reliance on mitochondrial DNA in previous studies [37], which can lead to overestimation of divergence times [60]. Consistent with previous studies, our Admixture analyses and range-wide effective migration surfaces suggest low gene flow between the PNW and Boreal lineages (Fig. 1C, Fig. 2A). The absence of observable gene flow is perhaps striking given their close proximity to each other in western North America. Melo-Ferreira et al. [37] hypothesized that these lineages may exhibit incipient reproductive isolation. The evolution of reproductive isolation between these populations is possible, but, given our more recent divergence time estimates, we suggest that reduced gene flow may reflect quite recent secondary contact between these populations in western North America. Genetic and fossil evidence suggests the common ancestors of PNW and Boreal populations occupied separate refugia in southern and eastern North America, respectively, during the last glacial maximum [36]. The east-to-west phylogenetic clustering (Fig. 1B) and genetic diversity gradient (Fig. 2B) of the Boreal population is consistent with a recent range expansion from this eastern refugia [36], which implies the PNW and Boreal hares have only recently experienced secondary contact. However, more detailed population history modeling and sampling of the putative contact zone is required to test these hypotheses.

Consistent with low historical gene flow, the AK winter-brown hare lacks the PNW winter-brown *Agouti* haplotype (Fig. 3), indicating a role for independent mutation leading to the convergent evolution of brown winter coats. This likely reflects constraints on the migration of adaptive variation between populations. Even in the absence of intrinsic or extrinsic reproductive isolation, the rate at which a positively selected variant spreads across a uniform environment is constrained by dispersal distance and the strength of positive selection [6]. Across spatially or temporally heterogenous landscapes, the spread of an adaptive allele between patches is further constrained by the strength or frequency of purifying selection in maladaptive habitats or climatic periods [2]. Given that previous models suggest a low probability of winter-brown camouflage along the majority of the Pacific coast [28], we suspect that the winter-brown PNW variant would have to traverse snowy habitats where it is strongly disfavored in order to reach Alaska, ∼3000 away via the coast. At this distance, the probability of adaptation through gene flow is virtually zero under a range of model assumptions (Fig. 4). In fact, dispersal limitations and patchy range-edge habitats should generally favor independent evolution of winter-brown camouflage at scales beyond 100-1000 km (depending on the assumed values of *μ* and *s*_*m*_; Fig. 4). Further research is needed to dissect the genetic basis of winter-brown camouflage in the northwestern and eastern edges of the range (e.g., Alaska and Pennsylvania). Subtle phenotypic similarities between observed winter-brown hares in Alaska and Pennsylvania could imply a shared genetic basis (e.g., white feet versus brown feet in the PNW, unpublished data). However, our theoretical modelling would suggest that they likely reflect independent mutations as well (i.e., ∼5900 km between AK and PA sampling sites).

Convergent evolution is thought to be more common for ‘loss-of-function’ traits because they may have a larger mutational target size relative to ‘gain-of-function’ traits [9,61]. Convergent color adaptation involving loss-of-function mutations has been shown between different species of lizard [9,62] and cavefish [63] and the repeated evolution of melanism across populations of deer mice involves convergent loss-of-function *Agouti* mutations [64]. In PNW hares, adaptive introgression of *Agouti* variation from black-tailed jackrabbits appears to have caused a reversion to the ancestral winter-brown condition in *Lepus* (i.e., the likely ancestral state before the common ancestor of winter-white *Lepus* species). As the ancestral winter-brown variant is recessive, this implies that derived winter-white camouflage in snowshoe hares evolved through dominant gain-of-function mutations, consistent with the seasonal upregulation of *Agouti* during the development of white coats [29]. Independent origins of winter-brown camouflage across the snowshoe hare range could similarly involve relatively simple loss-of-function mutations that break the molecular pathways contributing to the development of white winter coats. Indeed, the evolution of darkened winter coats in some populations of mountain hares (*Lepus timidus*) appears to have also involved introgression of a recessive, loss-of-function *Agouti* variant [59]. Collectively, our results suggest that geographic distance and mutational target size, in addition to population structure and history, should play a crucial role in generating hypotheses about the relative roles of independent mutation and gene flow in adaptation.

Despite the evidence for independent winter-brown mutations at broad spatial scales, we also find that the PNW *Agouti* allele has traversed a substantial distance (∼600 km) to western Montana – a predominately winter-white locality – where it occurs at a frequency of 1.24%. At this starting frequency, both theory and simulations show that a shift in environmental conditions towards favoring the brown allele results in a ∼80% probability of fixation (Fig. 5). The high probability of adaptation through gene flow ∼600 km from a focal patch is consistent with a scenario of very weak purifying selection (*s*_*m*_=0.0001) against the winter-brown allele in snowy environments under the Ralph and Coop [2] model (*P*(independent mutation)=∼3% (*μ*=4.54e-8) or ∼76% (*μ*=4.54e-6); Fig. 4). Although winter-brown camouflage is likely more strongly deleterious in winter-white environments given the known fitness consequences of mismatch [30], purifying selection against the winter-brown *Agouti* allele in winter-white environments may be weak or absent because it is recessive and thus hidden to selection at low frequency. In agreement with Ralph and Coop [2], we suggest that adaptation between distant populations via gene flow may be relatively more common through recessive variation compared to dominant variation when purifying selection is strong in intervening habitats.

Climate change is expected to reduce winter snow cover across the snowshoe hare range [26], which could potentially result in winter-brown camouflage being favored in predominately winter-white populations [28]. Under this scenario, positive selection could operate on convergent *de novo* winter-brown mutations or on standing genetic variation for winter-brown camouflage to facilitate rapid adaptation. Using simulations of positive selection, we show that although dominant and recessive mutations in mutation/migration-selection balance share a similar fixation probability (Fig. 5A), they are likely to experience vastly different frequency change dynamics (Fig. 5B) [24] and thus lead to very different rates of adaptation following environmental change. For instance, we show that in MT the initial rate of adaptation from segregating winter-brown *Agouti* variation is likely to be slow relative to an equivalently beneficial dominant mutation. While recessive mutations tended to fix more quickly than dominant mutations, they also took substantially longer to reach high frequencies in populations (Fig. 5B; e.g., 8837 vs. 1007 generations for a phenotype determined by recessive vs. dominant mutation to reach a population frequency of 75%, respectively). This pattern can be explained by the different allele frequency phases most strongly affected by genetic drift. For instance, positive selection can readily act on beneficial dominant mutations at low frequency, but at high frequency the recessive wildtype allele is hidden in heterozygotes and genetic drift governs (and generally slows) fixation dynamics. The opposite is true for beneficial recessive mutations, which are governed by genetic drift at low frequency but driven to fixation by selection. Overall, selection on low frequency recessive variation may be a relatively ineffective way to adapt rapidly to changing environments. In fact, the temporal lag for the spread of beneficial recessive variation may be sufficient enough to allow time for dominant independent mutations (e.g., gain-of-function MC1R mutations that result in melanism) [65] to appear and spread before the recessive allele increases appreciably. These findings highlight the important roles of genetic dominance in shaping both the geographic scope and rate of convergent adaptation and underscores the need for further study.

Collectively, our study provides important insights into long-standing evolutionary questions, as well as the potential for adaptation to climate change in snowshoe hares. A key to understanding the potential of adaptive variation to spread under climate change is revealing how it has spread in the past. We show that adaptive responses to reduced snow cover in snowshoe hares have involved both the local spread of winter-brown camouflage in the PNW as well as convergent evolution elsewhere in the range. In snowshoe hares and other seasonally changing species, regions of winter-camouflage polymorphism may be crucial areas to maintain population connectivity as a conduits for the spread of winter-brown camouflage across broader portions of the range [28]. However, facilitating gene flow alone may not be sufficient to facilitate rapid adaptive responses, given the broad range of snowshoe hares and the genetic architecture of winter-brown camouflage variation. Rather, we suspect that rapid climate change responses in winter-white populations distant from winter-brown habitat may also likely involve independent origins of winter-brown camouflage and selection on variation in coat color phenology (i.e., shifts in the timing of winter-white molts). Although the genetic basis of variation in camouflage phenology remains unresolved in this system, it is likely quantitative and perhaps more responsive to selection. Regardless, this study represents an important step towards making predictions about evolutionary responses under climate change in this species.

## Acknowledgements

We thank E. Cheng and K. Garrison for assistance with sample collection. We thank L. Olson for informing us on the recent collection of a winter brown snowshoe hare, and L. Olson and the University of Alaska Museum of the North facilitating a tissue loan of this specimen (UAM 116170). Our research would not be possible without the incredible and irreplaceable support of natural history museums. We thank J. Melo-Ferreira, P. C. Alves, M. S. Ferreira, N. Herrera, E. Kopania, A. Kumar, M. Zimova, K. Garrison, and the UNVEIL network for helpful discussions. Funding and support for this research was provided a National Science Foundation (NSF) Graduate Research Fellowship (DGE-1313190), NSF Doctoral Dissertation Improvement Grant (DGE-1702043), NSF Graduate Research Opportunities Worldwide, NSF EPSCoR (OIA-1736249), and NSF (DEB-0841884; DEB-1907022), the Drollinger-Dial Foundation, American Society of Mammalogists Grant-in-aid of Research, and a Swiss Government Excellence Scholarship.

## Data Accessibility

Original sequence data are available in the Sequence Read Archive (www.ncbi.nlm.nih.gov/sra). Previously generated whole exome and genome sequence data of snowshoe hare (BioProject PRJNA420081, SAMN02782769, SAMN07526959) are also available in the Sequence Read Archive.

## Author Contributions

MRJ, JMG, and LSM conceived and designed the study; LSM led all field work and related collection of biological samples; MRJ performed molecular lab work and data analysis, with contributions and guidance from JMG and JDJ; MRJ led the writing with contributions from all authors. All authors gave final approval for publication and agree to be held accountable for the work performed therein.

