## Supplemental Table for "Convergent evolution of seasonal camouflage in response to reduced snow cover across the snowshoe hare range"

**Table S1. Sample list and metadata**

| Sample ID | Symbol | State/Province | Latitude | Longitude | Sequencing |
| --- | --- | --- | --- | --- | --- |
| NBERG_67 | WY | Wyoming | 43.7614169 | -110.5074 | exome |
| !van114 | CO | Colorado | 38.9881268 | -106.73657 | exome |
| D_Olsen1 | UT | Utah | 39.8767395 | -110.80418 | exome,WGS |
| Gomm5 | ME | Maine | 45.035556 | -68.681944 | exome |
| Huston_13 | NY | New York | 43.952618 | -73.728266 | exome |
| Allab_3 | PA | Pennseylvania | 41.14675 | -75.58314 | exome,WGS |
| Mortensen_4 | MN | Minnesota | 47.7293567 | -94.734496 | exome |
| Nolson15 | AK2 | Alaska | 67.021483 | -162.44963 | exome |
| KKIEL32 | AK1 | Alaska | 64.6935 | -148.29169 | exome |
| Canada_213 | NS | Nova Scotia | 44.6592593 | -63.266104 | exome |
| Canada_193 | BC2 | British Columbia | 51.56681 | -121.32603 | exome |
| Canada_231 | YK | Yukon | 63.52549 | -136.20675 | exome |
| Canada_227 | BC3 | British Columbia | 59.38888 | -123.29424 | exome |
| Canada_221 | NT | Northwest Territories | 65.3 | -126.80008 | exome |
| Canada_53 | SK | Saskatchewan | 53.77753 | -106.91551 | exome |
| UAM_116170 | AK | Alaska | 58.69037 | -156.685 | WGS |
| STRAUSER53 | OR | Oregon | 44.4616623 | -121.97764 | exome |
| STRAUSER54 | OR | Oregon | 44.4625086 | -121.97763 | exome |
| STRAUSER55 | OR | Oregon | 44.4629421 | -121.97779 | exome |
| STRAUSER56 | OR | Oregon | 44.4627943 | -121.97637 | exome |
| STRAUSER57 | OR | Oregon | 44.4622388 | -121.97365 | exome |
| STRAUSER58 | OR | Oregon | 44.462774 | -121.9731 | exome |
| STRAUSER59 | OR | Oregon | 44.463033 | -121.97388 | exome |
| STRAUSER61 | OR | Oregon | 44.4601096 | -121.9822 | exome |
| STRAUSER62 | OR | Oregon | 44.4606075 | -121.9815 | exome |
| STRAUSER63 | OR | Oregon | 44.4173808 | -122.01279 | exome |
| STRAUSER64 | OR | Oregon | 44.4631437 | -121.97819 | exome |
| STRAUSER65 | OR | Oregon | 44.4627943 | -121.97637 | exome |
| STRAUSER66 | OR | Oregon | 44.4626872 | -121.97246 | exome |
| STRAUSER67 | OR | Oregon | 44.4630864 | -121.9738 | exome |
| STRAUSER68 | OR | Oregon | 44.4631437 | -121.97819 | exome |
| 8801 | OR | Oregon | 44.33119 | -109.97555 | exome |
| 8826 | OR | Oregon | 44.46121 | -109.97606 | exome |
| 8898 | OR | Oregon | 44.46131 | -109.97545 | exome |
| 8897 | OR | Oregon | 44.46164 | -109.97555 | exome,WGS |
| 8899 | OR | Oregon | 44.46163 | -109.97573 | exome |
| 8302 | OR | Oregon | 44.46158 | -109.97529 | exome |
| STRAUSER79 | OR | Oregon | 43.142354 | -121.70504 | exome |
| STRAUSER81 | OR | Oregon | 43.1413032 | -121.70131 | exome |
| STRAUSER82 | OR | Oregon | 43.1437813 | -121.70305 | exome |
| STRAUSER83 | OR | Oregon | 43.1413032 | -121.70131 | exome |
| STRAUSER84 | OR | Oregon | 43.1410209 | -121.70103 | exome |
| A0616 | WA | Washington | 47.681387 | -121.26308 | exome |
| A0951 | WA | Washington | 47.681387 | -121.26308 | exome |
| A0953 | WA | Washington | 47.681387 | -121.26308 | exome |
| A0952 | WA | Washington | 47.675159 | -121.27611 | exome |
| A0954 | WA | Washington | 47.675159 | -121.27611 | exome |
| STRAUSER34 | WA | Washington | 47.7971178 | -121.27668 | exome |
| STRAUSER35 | WA | Washington | 47.7980728 | -121.27497 | exome |
| STRAUSER36 | WA | Washington | 47.7966832 | -121.27531 | exome |
| STRAUSER37 | WA | Washington | 47.7986722 | -121.27654 | exome |
| STRAUSER38 | WA | Washington | 47.7969383 | -121.27372 | exome |
| STRAUSER39 | WA | Washington | 47.7967275 | -121.27346 | exome |
| STRAUSER40 | WA | Washington | 47.7998239 | -121.28494 | exome |
| STRAUSER41 | WA | Washington | 47.7986722 | -121.27654 | exome |
| STRAUSER42 | WA | Washington | 47.7967275 | -121.27346 | exome |
| STRAUSER43 | WA | Washington | 47.7970248 | -121.27289 | exome |
| 8875 | WA | Washington |  |  | exome |
| A0602 | WA | Washington | 47.675159 | -121.27611 | exome |
| A0903 | WA | Washington | 47.681387 | -121.26308 | exome |
| A0983 | WA | Washington | 47.675159 | -121.27611 | exome |
| A0991 | WA | Washington | 47.681387 | -121.26308 | exome |
| A0961 | WA | Washington | 47.681387 | -121.26308 | exome,WGS |
| A0965 | WA | Washington | 47.681387 | -121.26308 | exome |
| A0604 | WA | Washington | 47.681387 | -121.26308 | exome |
| A0603 | WA | Washington | 47.681387 | -121.26308 | exome |
| A0621 | WA | Washington | 47.681387 | -121.26308 | exome |
| A0901 | WA | Washington | 47.681387 | -121.26308 | exome |
| Woody1 | MT | Montana |  |  | exome |
| Woody2 | MT | Montana |  |  | exome |
| Woody3 | MT | Montana |  |  | exome |
| Woody4 | MT | Montana |  |  | exome |
| 8892 | MT | Montana |  |  | exome |
| 3651 | MT | Montana |  |  | exome |
| 8919 | MT | Montana |  |  | exome |
| 3680 | MT | Montana |  |  | exome |
| 6969 | MT | Montana |  |  | exome |
| 8150 | MT | Montana |  |  | exome |
| 8151 | MT | Montana |  |  | exome |
| 3032 | MT | Montana |  |  | exome |
| 3676 | MT | Montana |  |  | exome |
| 3686 | MT | Montana |  |  | exome |
| LAM.INE.2013 | MT | Montana |  |  | WGS |
| 08_1461 | BC1 | British Columbia | 49.16 | -122.58 | exome |
| 07_1119 | BC1 | British Columbia | 49.16 | -122.58 | exome |
| 08_2522 | BC1 | British Columbia | 49.28 | -122.79 | exome |
| 07_1665 | BC1 | British Columbia | 49.28 | -122.79 | exome |
| 07_2243 | BC1 | British Columbia |  |  | exome |
| 07_1618 | BC1 | British Columbia |  |  | exome |
| 06_1401 | BC1 | British Columbia | 49.16 | -122.58 | exome |
| 06_1209 | BC1 | British Columbia | 49.28 | -122.79 | exome |
| PZ001 | BC1 | British Columbia | 49.16 | -122.58 | exome |
| PZ002 | BC1 | British Columbia | 49.28 | -122.79 | exome |
| PZ005 | BC1 | British Columbia | 49.28 | -122.79 | exome |
| 12-0067 | BC1 | British Columbia |  |  | exome |
| 06_0627 | BC1 | British Columbia | 49.24243 | -122.94607 | exome,WGS |
| 06_0766 | BC1 | British Columbia | 49.16 | -122.58 | exome |
| MacKay2 | BTJR-NV1 | Nevada | 40.95705 | -115.46569 | WGS |
| MacKay3 | BTJR-NV2 | Nevada | 40.53012 | -115.33222 | WGS |
| Burke1 | BTJR-OR2 | Oregon | 43.372083 | -121.074 | WGS |
| Burke4 | BTJR-OR1 | Oregon | 43.372083 | -121.074 | WGS |
| Bauer6 | BTJR-CA | California | 40.912994 | -121.71212 | WGS |
| CHENG_R1834 | MT | Montana | 48.808515 | -114.28188 | pooled WGS |
| CHENG_R1851 | MT | Montana | 48.487527 | -113.36175 | pooled WGS |
| CHENG_R1986 | MT | Montana | 48.808515 | -114.28188 | pooled WGS |
| CHENG_R1988 | MT | Montana | 48.808515 | -114.28188 | pooled WGS |
| CHENG_R3000 | MT | Montana | 48.808515 | -114.28188 | pooled WGS |
| CHENG_R3005 | MT | Montana | 48.7765186 | -114.27609 | pooled WGS |
| CHENG_R3012 | MT | Montana | 48.3567919 | -113.31766 | pooled WGS |
| CHENG_R3013 | MT | Montana | 48.808515 | -114.28188 | pooled WGS |
| CHENG_R3019 | MT | Montana | 48.808515 | -114.28188 | pooled WGS |
| CHENG_R3027 | MT | Montana | 48.808515 | -114.28188 | pooled WGS |
| CHENG_R3056 | MT | Montana | 48.487527 | -113.36175 | pooled WGS |
| CHENG_R3057 | MT | Montana | 48.808515 | -114.28188 | pooled WGS |
| CHENG_R3058 | MT | Montana | 48.3567919 | -113.31766 | pooled WGS |
| CHENG_R3076 | MT | Montana | 48.49119 | -113.33836 | pooled WGS |
| CHENG_R3077 | MT | Montana | 48.49119 | -113.33836 | pooled WGS |
| CHENG_R3078 | MT | Montana | 48.808515 | -114.28188 | pooled WGS |
| CHENG_R3079 | MT | Montana | 48.8004028 | -113.70306 | pooled WGS |
| CHENG_R3080 | MT | Montana | 48.8004028 | -113.70306 | pooled WGS |
| CHENG_R3081 | MT | Montana | 48.8004028 | -113.70306 | pooled WGS |
| CHENG_R3083 | MT | Montana | 48.771661 | -114.26701 | pooled WGS |
| CHENG_R3084 | MT | Montana | 48.808515 | -114.28188 | pooled WGS |
| CHENG_R3096 | MT | Montana | 48.808515 | -114.28188 | pooled WGS |
| CHENG_R3098 | MT | Montana | 48.808515 | -114.28188 | pooled WGS |
| CHENG_R3354 | MT | Montana | 48.3567919 | -113.31766 | pooled WGS |
| CHENG_R3356 | MT | Montana | 48.808515 | -114.28188 | pooled WGS |
| CHENG_R3358 | MT | Montana | 48.808515 | -114.28188 | pooled WGS |
| CHENG_R3359 | MT | Montana | 48.808515 | -114.28188 | pooled WGS |
| CHENG_R3360 | MT | Montana | 48.808515 | -114.28188 | pooled WGS |
| CHENG_R3375 | MT | Montana | 48.49119 | -113.33836 | pooled WGS |
| CHENG_R3376 | MT | Montana | 48.49119 | -113.33836 | pooled WGS |
| CHENG_R3390 | MT | Montana | 48.808515 | -114.28188 | pooled WGS |
| CHENG_R3413 | MT | Montana | 48.49119 | -113.33836 | pooled WGS |
| CHENG_R3432 | MT | Montana | 48.49119 | -113.33836 | pooled WGS |
| CHENG_R3434 | MT | Montana | 48.808515 | -114.28188 | pooled WGS |
| CHENG_R3436 | MT | Montana | 48.808515 | -114.28188 | pooled WGS |
| CHENG_R3437 | MT | Montana | 48.808515 | -114.28188 | pooled WGS |
| CHENG_R3438 | MT | Montana | 48.3567919 | -113.31766 | pooled WGS |
| CHENG_R3441 | MT | Montana | 48.808515 | -114.28188 | pooled WGS |
| CHENG_R3443 | MT | Montana | 48.808515 | -114.28188 | pooled WGS |
| CHENG_R3446 | MT | Montana | 48.808515 | -114.28188 | pooled WGS |
| CHENG_R3449 | MT | Montana | 48.808515 | -114.28188 | pooled WGS |
| CHENG_R3478 | MT | Montana | 48.487527 | -113.36175 | pooled WGS |
| CHENG_R3485 | MT | Montana | 48.8004028 | -113.70306 | pooled WGS |
| CHENG_R3489 | MT | Montana | 48.8004028 | -113.70306 | pooled WGS |
| CHENG_R3500 | MT | Montana | 48.49119 | -113.33836 | pooled WGS |
| CHENG_R3503A | MT | Montana | 48.3567919 | -113.31766 | pooled WGS |
| CHENG_R3504 | MT | Montana | 48.808515 | -114.28188 | pooled WGS |
| CHENG_R3506 | MT | Montana | 48.487527 | -113.36175 | pooled WGS |
| CHENG_R3511 | MT | Montana | 48.808515 | -114.28188 | pooled WGS |
| CHENG_R3517 | MT | Montana | 48.808515 | -114.28188 | pooled WGS |
| CHENG_R3574 | MT | Montana | 48.487527 | -113.36175 | pooled WGS |
| CHENG_R3586 | MT | Montana | 48.49119 | -113.33836 | pooled WGS |
| CHENG_R3587 | MT | Montana | 48.8004028 | -113.70306 | pooled WGS |
| CHENG_R3792 | MT | Montana | 48.8004028 | -113.70306 | pooled WGS |
| CHENG_R3882 | MT | Montana | 48.49119 | -113.33836 | pooled WGS |
| CHENG_R3896 | MT | Montana | 48.808515 | -114.28188 | pooled WGS |
| CHENG_R3897 | MT | Montana | 48.49119 | -113.33836 | pooled WGS |
| HARE105 | MT | Montana | 48.808515 | -114.28188 | pooled WGS |
| MILLS_R1483 | MT | Montana | 48.4565404 | -114.87639 | pooled WGS |
| MILLS_R1485 | MT | Montana | 48.4565404 | -114.87639 | pooled WGS |
| MILLS_R1487 | MT | Montana | 48.4565404 | -114.87639 | pooled WGS |
| MILLS_R1492 | MT | Montana | 48.4565404 | -114.87639 | pooled WGS |
| MILLS_R1501 | MT | Montana | 48.5472782 | -114.63168 | pooled WGS |
| MILLS_R1522 | MT | Montana | 48.5472782 | -114.63168 | pooled WGS |
| MILLS_R1524 | MT | Montana | 48.5472782 | -114.63168 | pooled WGS |
| MILLS_R1744 | MT | Montana | 48.4565404 | -114.87639 | pooled WGS |
| MILLS_R7189 | MT | Montana | 48.5472782 | -114.63168 | pooled WGS |
| MILLS_R7307 | MT | Montana | 48.4565404 | -114.87639 | pooled WGS |
| MILLS_R7327 | MT | Montana | 48.4565404 | -114.87639 | pooled WGS |
| MILLS_R7425 | MT | Montana | 47.4086908 | -113.55456 | pooled WGS |
| MILLS_R7544 | MT | Montana | 47.4086908 | -113.55456 | pooled WGS |
| MILLS_R7550 | MT | Montana | 47.4086908 | -113.55456 | pooled WGS |
| MILLS_R7638 | MT | Montana | 47.148166 | -113.73541 | pooled WGS |
| MILLS_R7707 | MT | Montana | 47.148166 | -113.73541 | pooled WGS |
| MILLS_R7712 | MT | Montana | 47.148166 | -113.73541 | pooled WGS |
| MILLS_R7723 | MT | Montana | 47.148166 | -113.73541 | pooled WGS |
| MILLS_R7749 | MT | Montana | 47.148166 | -113.73541 | pooled WGS |
| MILLS_R7809 | MT | Montana | 47.148166 | -113.73541 | pooled WGS |
| MILLS_R7819 | MT | Montana | 47.4086908 | -113.55456 | pooled WGS |
| MILLS_R7920 | MT | Montana | 47.148166 | -113.73541 | pooled WGS |
| MILLS_R8238 | MT | Montana | 48.4565404 | -114.87639 | pooled WGS |
